# Distinguishing matrix-associated mineralization from solution-driven calcium phosphate precipitation in osteogenic culture

**DOI:** 10.64898/2026.10.27.762140

**Authors:** Ali Aslan Demir, Thomas Combriat

**Affiliations:** NJORD Centre, University of Oslo, Oslo, Norway

**Keywords:** osteogenic culture, *β*-glycerophosphate, calcium phosphate, kinetic modelling, mineralization

## Abstract

Mineral accumulation in osteogenic culture is often interpreted as evidence of cell-guided extracellular matrix mineralization. However, measurements could also capture calcium-phosphate precipitation outside of the organized extracellular matrix. High concentrations of *β*-glycerophosphate (*β*-GP) could promote nonphysiological mineral deposition. We developed a kinetic model to examine when mineral deposition occurs within the matrix and when precipitation from the medium becomes the dominant mechanism.

The model couples *β*-GP hydrolysis, medium renewal, matrix-associated precursor formation and maturation, and solution-driven precipitation. We examined model behavior using parameter sweeps, global sensitivity analysis, comparison with published mineralization time courses, and equifinality analysis to determine whether similar mineralization amounts could arise from different pathway contributions and parameter combinations.

Under baseline conditions (10 mM *β*-GP, 2-day renewal), the matrix-associated pathway activated first, followed by solution-driven precipitation about 18 h later. Once matrix capacity was approached, additional phosphate mainly increased non-matrix deposition. Parameter sets with similar total mineral gave matrix-associated fractions ranging from 0.17 to 1.00. Thus, the total amount of mineral does not necessarily indicate how much formed within the matrix, highlighting the importance of phosphate supply and medium renewal in interpreting osteogenic culture results.

## 1 Introduction

Bone mineralization is a coupled biological and physicochemical process regulated within the extracellular matrix [1, 2]. Osteoblasts organize this environment through matrix production, local control of calcium and phosphate ions, and mineralization-associated proteins [3, 4, 5]. Mineral formation depends not only on calcium and phosphate availability, but also on where these ions accumulate and how the matrix supports nucleation and growth [6]. Reproducing the complex organization is important for in vitro osteogenic models and bone tissue-engineering systems [7, 8].

Within the bone extracellular matrix, mineral formation is closely associated with matrix vesicles and collagen. Matrix vesicles concentrate calcium and phosphate and provide early sites for mineral formation, while collagen supports later mineral growth and organization [9, 10]. Several studies indicate that mineral formation can pass through a transient amorphous calcium phosphate (ACP) phase, a metastable and poorly ordered precursor that can subsequently transform toward more crystalline apatite-like mineral [9, 11, 6, 12, 13, 14, 15]. During the maturation step, mineralization becomes poorly crystalline, carbonated, ion-substituted nanoscale apatite rather than stoichiometric hydroxyapatite [11, 16, 14, 15].

In osteogenic culture, *β*-GP increases soluble phosphate through alkaline-phosphatase-mediated hydrolysis [17, 18, 19, 20]. The released phosphate can support matrix mineralization, but at higher levels it can also promote calcium-phosphate precipitation outside the organized extracellular matrix [19, 21, 22, 23]. Calcium-phosphate mineral can therefore arise through biologically distinct routes, depending on where nucleation occurs and how the local matrix environment facilitates mineral formation [13, 24, 6]. Conventional mineralization assays do not resolve these contributions, so a stronger endpoint signal may reflect greater matrix-associated mineralization, greater solution-driven precipitation, or both [25, 23]. Bonewald et al. showed that positive von Kossa staining did not necessarily correspond to bone-like apatite formation, highlighting the need for independent mineral characterization [26]. This raises a practical question for in vitro osteogenic culture: under what conditions does a bulk mineralization endpoint primarily reflect matrix-associated mineralization, and when might solution-driven precipitation contribute substantially to the same signal?

Experimental studies have reported acellular *β*-GP mineralization [27, 21] and dose-dependent non-specific deposition [28], further illustrating the difficulty of interpreting bulk mineral accumulation. Kinetic modeling offers a way to examine how different deposition routes may contribute under changing culture conditions [29]. Previous physicochemical models have described hydroxyapatite growth using semi-empirical expressions based on supersaturation, ion concentrations, and reactive surface area [30]. However, rate constants obtained from crystal growth experiments in solution are not directly applicable to osteogenic cultures. Cells and extracellular matrix provide additional nucleation sites and different growth surfaces, while local ion regulation can further affect mineral formation rates [31]. Matrix mineralization and bone formation have also been modeled [32], but the competition between matrix-associated formation and solution-driven precipitation within the same osteogenic culture remains less explored.

We developed an in vitro culture-scale kinetic model coupling *β*-GP hydrolysis and periodic medium renewal to two competing routes: matrix-associated precursor formation and maturation, and solution-driven calcium-phosphate precipitation. Both draw from the same soluble calcium and phosphate pools. The model examines how phosphate supply, renewal interval, and finite matrix capacity affect their relative contributions. It also explores how similar bulk mineralization endpoints can arise from different deposition pathways, pointing to conditions that could be tested experimentally.

## 2 Model

### 2.1 Model scope and rationale

The model describes calcium-phosphate accumulation in a well-mixed osteogenic culture with periodic medium renewal. Soluble calcium, phosphate, and *β*-GP are replenished, while formed mineral is retained. Two pathways are considered: matrix-associated formation through an ACP-like precursor that matures into apatite-like mineral (A and H), and solution-driven precipitation (Q) at a higher driving force.

Mineral origin is defined by the model compartments, with pathway contributions depending on the assumed kinetics, onset thresholds, and retention capacities. We used a phenomenological description rather than explicitly modelling individual cellular processes. Spatial transport and aqueous speciation are accounted for through effective parameters, avoiding further mechanistic detail that would be difficult to constrain from available data. Matrix vesicles, collagen, and ECM organization are important for mineral formation, but modeling each process would require additional mechanisms and experimental constraints. Their influence is incorporated into the matrix-associated pathway, which assumes preferential access to mineralization sites. Moreover, bulk mineralization measurements cannot distinguish between the two deposition routes without a set of controlled experiments, leaving additional mechanistic parameters poorly constrained. We therefore focused on the main processes governing their competition rather than describing mineral formation in greater cellular detail.

The two mineralization routes are assigned separate activation thresholds, representing differences in their effective nucleation conditions. Matrix-associated precursor formation is assumed to be accessible at a lower driving force than precipitation in the surrounding medium. The physical basis and limitations of this assumption are discussed in Section 2.3.

The lower threshold therefore represents matrix-facilitated ACP-like precursor formation in biological settings, while the higher effective threshold represents the onset of solution-driven precipitation from the surrounding medium. The matrix-associated route is described by the retained pools *A* and *H*, where *A* is the ACP-like precursor and *H* is the apatite-like mineral formed by its maturation. The solution-driven route is represented by the retained apatite-like pool *Q*.

The overall model structure is summarized in Fig. 1. Medium renewal restores the soluble species, while all mineral pools are accumulated over time.

**Figure 1.**
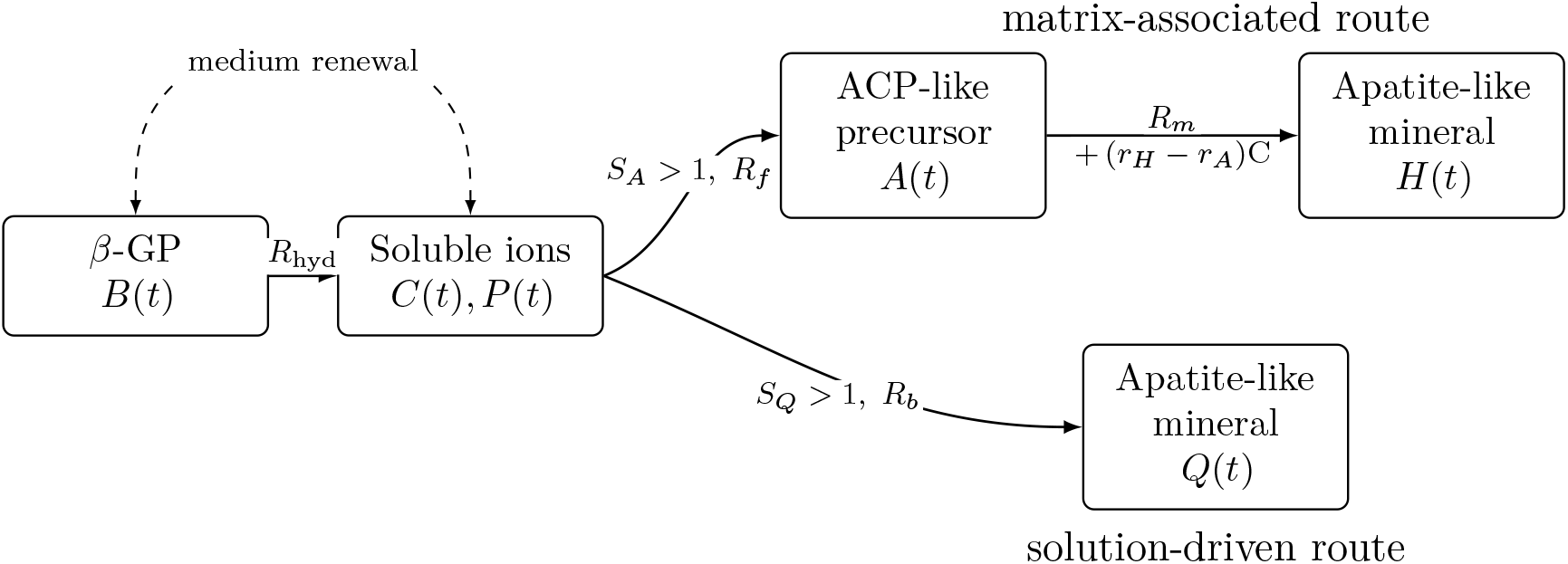
Schematic of the competing-pathway model. Hydrolysis of *β*-GP supplies soluble phosphate between medium renewals. The matrix-associated route becomes active when *S*_*A*_ *>* 1, forming an ACP-like precursor *A*(*t*) that matures into apatite-like mineral *H*(*t*). Because *K*_*Q*_ *> K*_*A*_, solution-driven precipitation requires a higher calcium-phosphate driving force and begins only when *S*_*Q*_ *>* 1, forming *Q*(*t*). All mineral pools are retained.

### 2.2 Competing mineralization pathways

The model describes phosphate release from *β*-GP and two competing mineralization routes, summarized in Eq. 1:

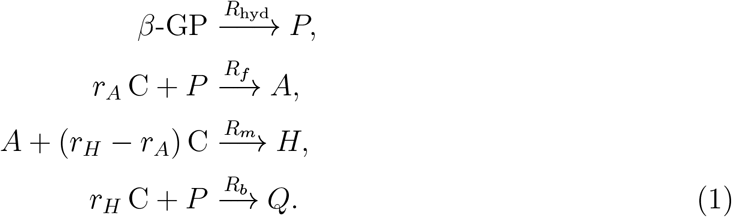

Here, *P* denotes soluble inorganic phosphate and *C* the soluble calcium concentration. *R*_hyd_ is the rate of *β*-GP hydrolysis. The matrix-associated pathway forms the ACP-like precursor *A* at rate *R*_*f*_, which then matures into the apatite-like pool *H* at rate *R*_*m*_. The solution-driven pathway forms *Q* at rate *R*_*b*_. Thus, *A* and *H* represent matrix-associated mineral, whereas *Q* represents mineral formed from precipitation in the surrounding solution.

The mineral pools are expressed on a phosphate-equivalent basis. The parameters *r*_*A*_ and *r*_*H*_ are the Ca/P ratios of the ACP-like and apatite-like pools, respectively. Since *r*_*H*_ *> r*_*A*_, conversion of *A* to *H* requires an additional *r*_*H*_ *− r*_*A*_ calcium equivalents per phosphate-equivalent unit [33]. The same value of *r*_*H*_ is used for both *H* and *Q*, because both are treated as apatite-like mineral.

Both pathways consume calcium and phosphate from the same soluble pool. Representing *Q* directly as an apatite-like pool is a culture-scale simplification and does not exclude transient amorphous phases during precipitation.

### 2.3 Formation indices and pathway activation

The formation indices *S*_*A*_ and *S*_*Q*_ describe the onset of the matrix-associated and solution-driven pathways, respectively:

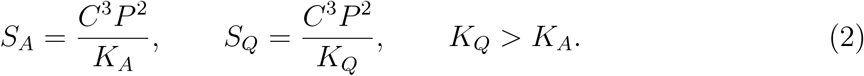

Here, *C* and *P* are the soluble calcium and phosphate concentrations. The parameters *K*_*A*_ and *K*_*Q*_, with units of mM^5^, define the onset conditions.

The ordering *K*_*A*_ *< K*_*Q*_ represents preferential nucleation within the matrix-associated environment. Matrix vesicles and collagen-associated sites can concentrate ions and facilitate nucleation, allowing matrix-associated mineralization at a lower calcium-phosphate driving force than precipitation in the surrounding medium [1, 2, 34, 24]. This is represented by *K*_*A*_ *< K*_*Q*_, although the actual onset times depend on phosphate release, calcium availability, and medium renewal. Marsh et al. also showed that mature osteoblast-derived extracellular matrix could support mineralization at lower calcium and phosphate availability by providing favorable nucleation sites [35].

Both thresholds are effective culture-scale parameters, not equilibrium solubility products [13, 9]. The formation indices do not explicitly account for ionic activities, phosphate speciation, pH, carbonate, complexation, or protein binding. Thus, *S*_*A*_ *>* 1 with *S*_*Q*_ *≤* 1 defines a matrix-selective regime, whereas *S*_*Q*_ *>* 1 permits solution-driven precipitation. Reported extracellular phosphate concentrations provide biological context rather than fixed activation thresholds [36].

### 2.4 Phosphate release

Hydrolysis of *β*-GP supplies soluble inorganic phosphate during osteogenic culture. *β*-GP can be hydrolysed by osteoblast-associated alkaline phosphatase and, depending on the medium, by serum-derived alkaline phosphatase [19, 18, 21, 20].

The hydrolysis rate is defined in Eq. 3 as

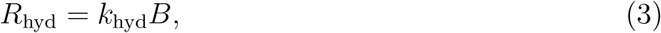

where *B* is the concentration of unhydrolysed *β*-GP and *k*_hyd_ is a hydrolysis rate constant. Within a renewal cycle beginning at time *t*_*j*_, let *τ* = *t − t*_*j*_, with 0 *≤ τ < T*_media_. Complete medium renewal sets 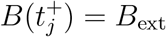, so Eq. 3 gives

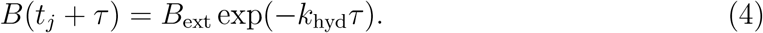

The fraction hydrolysed over a reference interval *T* is therefore *η*_*T*_ = 1 *−* exp(*−k*_hyd_*T* ), which gives

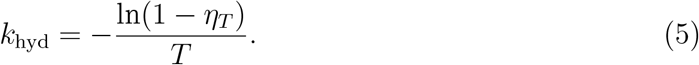

The baseline value of *k*_hyd_ was obtained from this relation by assuming that a fraction *η*_48*h*_ = 0.30 of *β*-GP is hydrolysed over 48 h. The literature supports ALP-mediated *β*-GP hydrolysis and variability in phosphate release across osteogenic culture conditions [20, 22].

The released phosphate increases *P* and raises both formation indices in Eq. 2. As phosphate accumulates between medium renewals, the system can shift from a matrix-selective regime to one in which solution-driven precipitation is also active. Rapid *β*-GP hydrolysis and elevated extracellular phosphate have been associated with nonphysiological calcium-phosphate deposition [19, 18, 21].

### 2.5 Mineralization rates and pathway competition

The matrix-associated precursor-formation rate, *R*_*f*_, is written as

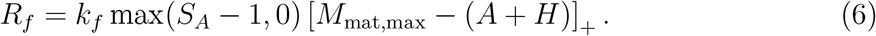

The parameter *k*_*f*_ is the matrix-associated formation coefficient, including the effects of cell activity and available matrix sites. Formation occurs only when the matrix-associated index exceeds its onset condition, *S*_*A*_ *>* 1, and the rate then increases with the excess driving force *S*_*A*_ *−* 1. The second factor limits further formation as the retained matrix-associated mineral, *A* + *H*, approaches the capacity *M*_mat,max_. At the culture scale, *k*_*f*_ therefore collects several effects that are not represented separately, including the abundance of mineralization-competent sites, the local matrix environment, and cell-dependent mineralization activity [10].

Solution-driven precipitation is described by

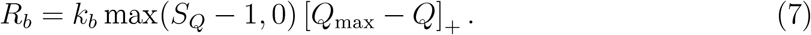

Here, *k*_*b*_ is the solution-driven formation coefficient. The rate remains zero until *S*_*Q*_ *>* 1. Once this threshold is crossed, precipitation is scaled by the remaining retention capacity, *Q*_max_ *− Q*, where *Q*_max_ is the maximum retained solution-origin mineral represented by the model. It is a culture-scale capacity, not a thermodynamic solubility limit.

The two rate laws therefore have the same general structure but act on different pools and different onset conditions. *M*_mat,max_ limits the cumulative matrix-associated pool *A* + *H*, whereas *Q*_max_ limits the cumulative solution-driven pool *Q*. These capacities apply over repeated medium-renewal cycles. Within each cycle, the amount of mineral that can form is still restricted by the soluble calcium and phosphate available through the elemental balances.

### 2.6 Precursor maturation

The matrix-associated precursor matures according to *R*_*m*_ = *k*_*m*_*A*, where *k*_*m*_ is the maturation coefficient in day^*−*1^. This first-order form is consistent with kinetic models in which ACP-to-apatite conversion is proportional to the amount of ACP present [37].

Reported conversion rates vary strongly with solution composition. Accordingly, *k*_*m*_ is treated as a culture-scale parameter rather than transferred directly from solution experiments. The term represents net conversion of *A* into *H* and does not distinguish between direct structural transformation and dissolution-reprecipitation [12, 13, 24, 6].

### 2.7 Medium renewal

Medium renewal is applied every *T*_media_. At each renewal event, *B, C*, and *P* are reset to *B*_ext_, *C*_ext_, and *P*_ext_, respectively. The mineral pools *A, H*, and *Q* are retained.

This assumes that solution-origin deposit stays with the culture when the medium is changed. Precipitate that is washed out with the medium is not modeled.

### 2.8 Governing equations

Between medium renewals, the model is described by

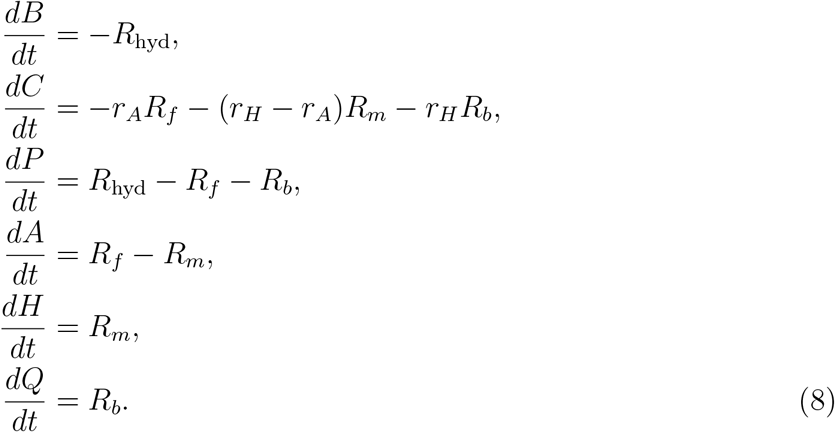

The first three equations describe changes in the soluble pools. *β*-GP is depleted by hydrolysis, phosphate is released and consumed by both mineralization pathways, and calcium is consumed according to the Ca/P ratio of each mineral-forming step.

The remaining equations track retained mineral. The precursor pool *A* increases through matrix-associated formation and decreases by maturation, while *H* increases through *R*_*m*_ and *Q* through *R*_*b*_. The rate expressions are given by Eqs. 3, 6, and 7, with *R*_*m*_ = *k*_*m*_*A*.

These balances conserve calcium and phosphate between media renewals.

*K*_*A*_ represents the lower onset of mineral formation supported by matrix vesicles and collagen, while *K*_*Q*_ represents precipitation outside the organized matrix. The coefficient *k*_*f*_ includes the effects of cell activity and available matrix sites, *k*_*m*_ describes precursor maturation, and *k*_*b*_ controls solution-driven precipitation. *M*_mat,max_ represents the mineral capacity of the matrix-associated pathway, while *Q*_max_ limits the amount of solution-origin mineral retained in the culture. These are culture-scale parameters, not direct measurements of cell activity or matrix properties.

### 2.9 Medium composition and model initialization

Simulations were initialized using ion concentrations representative of osteogenic *α*-MEM supplemented with *β*-GP. Basal calcium and phosphate concentrations were taken from the medium formulation given in Table 1.

**Table 1.** Medium composition and renewal conditions used for model initialization. Basal ion concentrations correspond to Gibco *α*-MEM [38]. The inorganic phosphate contribution from 10% FBS was estimated from reported commercial FBS composition [39].

| Component | Value | Model representation |
| --- | --- | --- |
| Calcium chloride (CaCl<sub>2</sub>) | 1.80 mM | Basal α-MEM calcium concentration. |
| Total calcium (10% FBS) | 2.00 mM | Estimated volume-weighted calcium concentration, $C_{\text{ext}} = 0.90(1.80) + 0.10(3.60) = 1.98 \text{ mM} \approx 2.00 \text{ mM}$ [40]. |
| Total phosphate, α-MEM | 1.014 mM | Basal phosphate concentration in α-MEM. |
| Total inorganic phosphate | 1.25 mM | Simulation input, $P_{\text{ext}} = 1.25 \text{ mM}$; estimated from $0.90(1.014) + 0.10(3.26) = 1.24 \text{ mM}$. |
| β-GP | 10.0 mM | Phosphate precursor, $B_{\text{ext}}$. |
| Medium-renewal interval | 2.0 d | Renewal period, $T_{\text{media}}$. |

The initial conditions are defined in Eq. 9 as

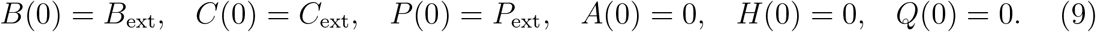

For the baseline simulation, the medium was replaced completely every 2 days.

## 3 Parameterization and Implementation

### 3.1 Baseline parameters and parameter selection

The baseline parameters were selected to represent a reference osteogenic culture, using reported medium compositions and literature evidence where available. A lower nucleation barrier was assumed for matrix-associated mineralization. The kinetic coefficients and retention capacities were assigned common reference values to avoid favouring either pathway initially. These parameters are not directly calibrated to a specific culture system. Their influence on the predictions was examined through parameter sweeps, global sensitivity analysis, and endpoint-equifinality analysis. Table 2 summarizes the kinetic, compositional, and empirical model parameters used for the reference simulations.

**Table 2.**
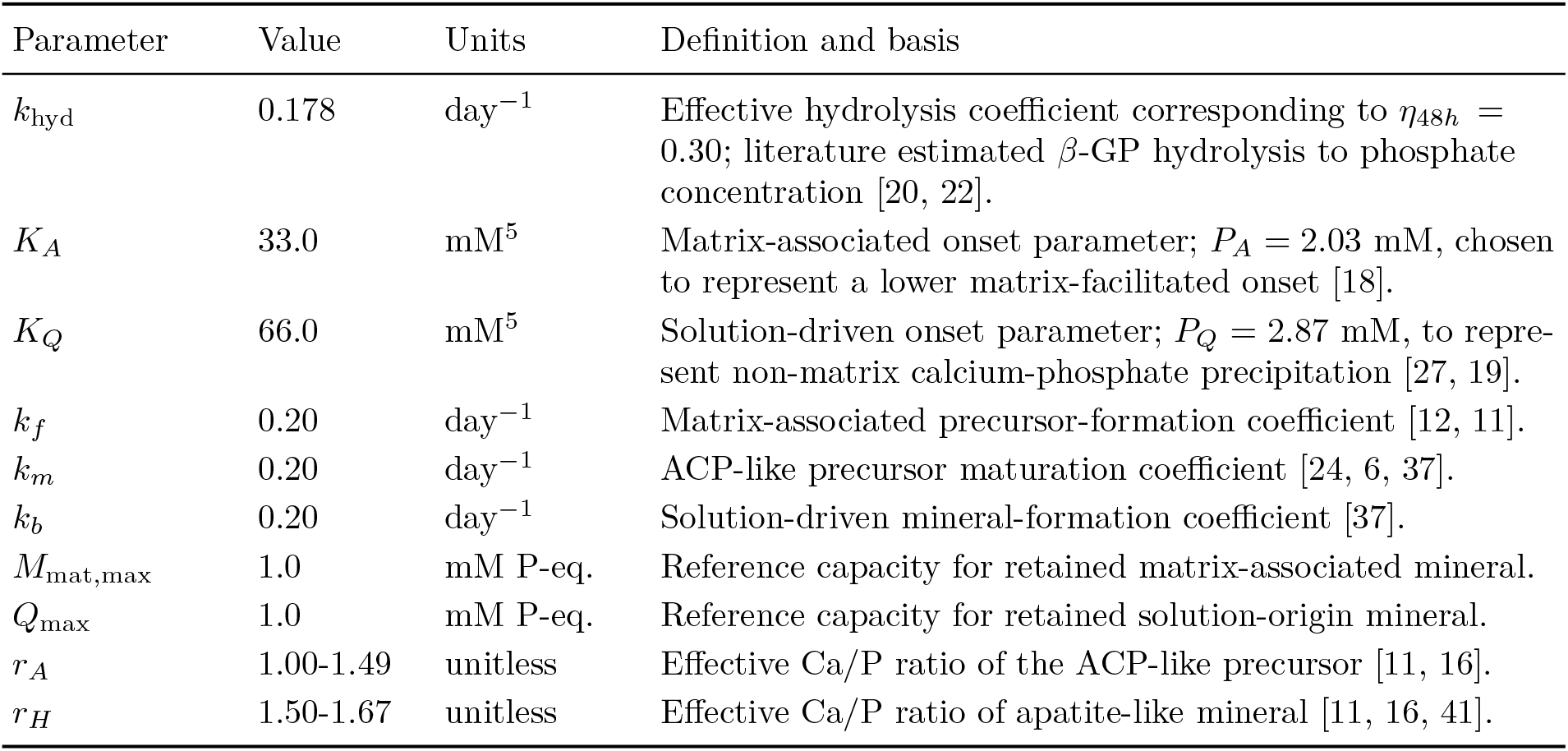
Baseline model parameters used in the simulations.

The onset parameters were expressed in terms of the phosphate concentration required for pathway activation, 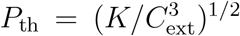. At *C*_ext_ = 2.00 mM, the baseline values *K*_*A*_ = 33.0 mM^5^ and *K*_*Q*_ = 66.0 mM^5^ correspond to *P*_*A*_ = 2.03 mM and *P*_*Q*_ = 2.87 mM, respectively. These values were treated as effective culture-scale onset parameters rather than thermodynamic solubility products or directly measured activation thresholds. The lower threshold represents matrix-facilitated formation of an ACP-like precursor, consistent with observations that elevated inorganic phosphate can initiate mineralization preferentially in osteoid-containing cultures [18]. Bellows et al. further showed that mineralization occurred in nodule-containing cultures but not in cultures lacking mineralizable osteoid, supporting a lower effective nucleation barrier for the matrix-associated pathway. The higher threshold represents the greater bulk driving force required for solution-driven calcium-phosphate deposition. This distinction is supported by studies showing that extensive hydrolysis of *β*-GP can strongly increase extracellular phosphate and promote calcium-phosphate deposition outside of a mineralizable osteoid context [27, 19]. Thus, the literature supports the ordering *K*_*A*_ *< K*_*Q*_ more directly than the exact numerical value of either onset parameter.

The kinetic coefficients *k*_*f*_, *k*_*m*_, and *k*_*b*_ were assigned a common baseline value of 0.20 day^*−*1^. For the first-order maturation term, this corresponds to a characteristic timescale of approximately 5 days. Equal values of *k*_*f*_ and *k*_*b*_ were used as a reference choice, although *K*_*A*_ *< K*_*Q*_ favors matrix-associated formation. These coefficients are treated as effective reference parameters rather than directly measured rate constants. Experimental studies support the sequence represented by the model, in which amorphous or highly disordered calcium-phosphate phases can precede more crystalline apatite-like mineral [12, 11, 6]. The first-order maturation term *R*_*m*_ = *k*_*m*_*A* is additionally consistent with kinetic descriptions in which ACP-to-apatite conversion is proportional to the amount of ACP present [37].

The capacity parameters *M*_mat,max_ and *Q*_max_ were both set to 1.0 mM P-eq. as common reference retention capacities. These values were not taken from direct experimental measurements and were chosen to avoid assigning a larger baseline capacity to either pathway. Their influence on model behavior was therefore evaluated separately in the parameter-sweep and global sensitivity analyses.

### 3.2 Numerical implementation

The model was implemented in Python using NumPy 2.4.4 and SciPy 1.17.1. The equations were integrated over 28 days using the adaptive RK45 solver (relative tolerance 10^*−*6^, absolute tolerance 10^*−*9^). Integration was restarted at each medium renewal, resetting soluble species while retaining mineral pools. Both the output interval and maximum solver step were set to Δ*t* = 0.05 day.

At each rate evaluation, mineralization rates were scaled down if the estimated calcium or phosphate consumption over Δ*t* exceeded the available amount. Rates requiring depleted ions were set to zero, while small negative recorded values were clipped to zero. Numerical step-size sensitivity was evaluated over Δ*t* = 0.0125–0.10 day, with negligible differences in mineralization trajectories and day-28 mineral amounts. The ion-availability limiter was not activated in the baseline simulations or in an additional screening of 50 parameter combinations. The convergence analysis and screening results are provided in the Supplementary Information.

Global sensitivity was assessed using nine independently and uniformly sampled parameters (in Supplementary). Scrambled Sobol sampling (*N* = 1024, seed 42), without second-order indices, resulted in 11,264 model evaluations. First-order and total-order indices were calculated for *A*(28) + *H*(28) and *Q*(28). Confidence intervals (95%) were estimated from 100 bootstrap resamples without a fixed bootstrap seed.

Endpoint equifinality was examined using 2000 Latin-hypercube samples in log-parameter space. The 50 samples closest to the reference endpoint were further refined by bounded optimization. Parameter sets within 2% of the reference day-28 total mineral were accepted.

Parameter ranges, sampling procedures, and verification results are reported in the Supplementary Information.

## 4 Results

### 4.1 Dynamics of matrix-associated and solution-driven pathway

Under the baseline parameters, soluble phosphate increased within each medium-renewal cycle as *β*-GP was hydrolysed, whereas soluble calcium was gradually depleted by mineral formation (Fig. 2A). The matrix-associated activation crossed its threshold early in the simulation. In the baseline scenario, the first numerical crossing of *S*_*A*_ = 1 occurred at 11 h, whereas *S*_*Q*_ = 1 was first crossed at 29 h (Fig. 2B).

**Figure 2.**
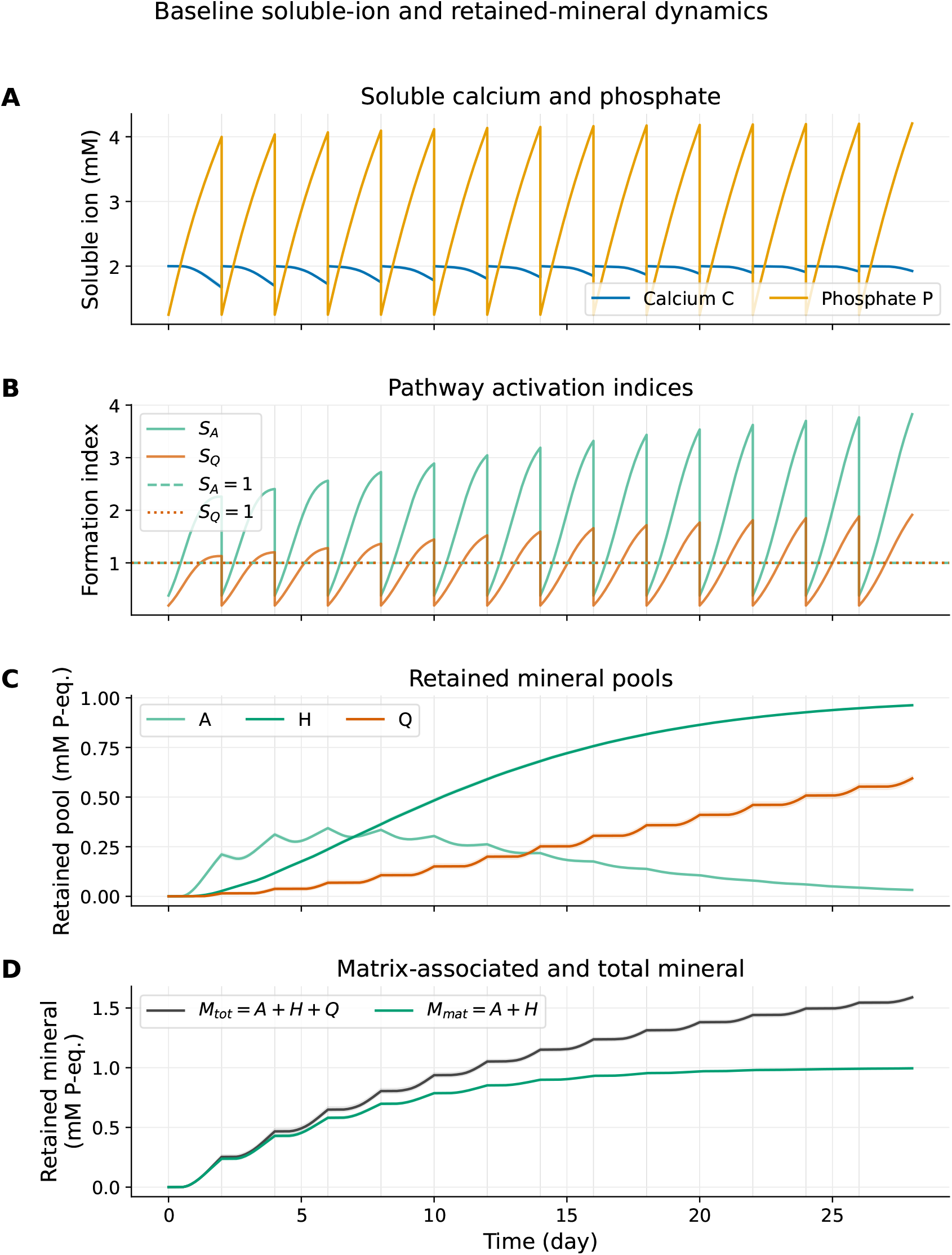
Baseline soluble-ion and mineral dynamics. **A**, Soluble calcium and phosphate during repeated medium-renewal cycles. **B**, Pathway activation indices *S*_*A*_ and *S*_*Q*_. **C**, Retained ACP-like precursor *A*, matrix-associated apatite-like mineral *H*, and solution-driven mineral *Q*. **D**, Matrix-associated mineral *M*_mat_ = *A* + *H* and total retained mineral *M*_tot_ = *A* + *H* + *Q*.

The retained pools showed the sequence of precursor formation and maturation. The ACP-like pool *A* increased first, reached a peak of 0.34 mM P-eq. at day 6, and then declined as the apatite-like matrix pool *H* accumulated (Fig. 2C). By day 28, *A* ≈ mM P-eq., *H* ≈ 0.96 mM P-eq., and *Q* ≈ 0.59 mM P-eq.. Thus, *M*_mat_ = *A* + *H* reached ≈ 0.99 mM P-eq., near the maximal matrix-associated capacity, while total retained mineral reached *M*_tot_ = *A* + *H* + *Q* ≈ 1.59 mM P-eq.. The day-28 matrix-associated fraction was ≈ 0.63.

Sampling *r*_*A*_ and *r*_*H*_ over their prescribed ranges caused little variation in matrix-associated mineral but modest variation in solution-driven mineral. The 95% interval for final *M*_mat_ was 0.99-1.00 mM P-eq., whereas final *Q* varied from 0.58 to 0.61 mM P-eq..

### 4.2 Medium renewal and phosphate dynamics govern pathway activation

Figure 3A presents an analytical approximation of pathway accessibility across *β*-GP concentrations and medium-renewal intervals. The estimated threshold crossings hold calcium fixed at *C*_ext_ and neglect phosphate consumption by mineralization. Greater phosphate supply or longer renewal intervals favor sequential access to the matrix-associated and solution-driven pathways. In the coupled model, however, mineral formation depletes soluble ions and may delay or prevent solution-driven activation. Thus, analytical accessibility does not necessarily imply pathway activation in the full simulation.

**Figure 3.**
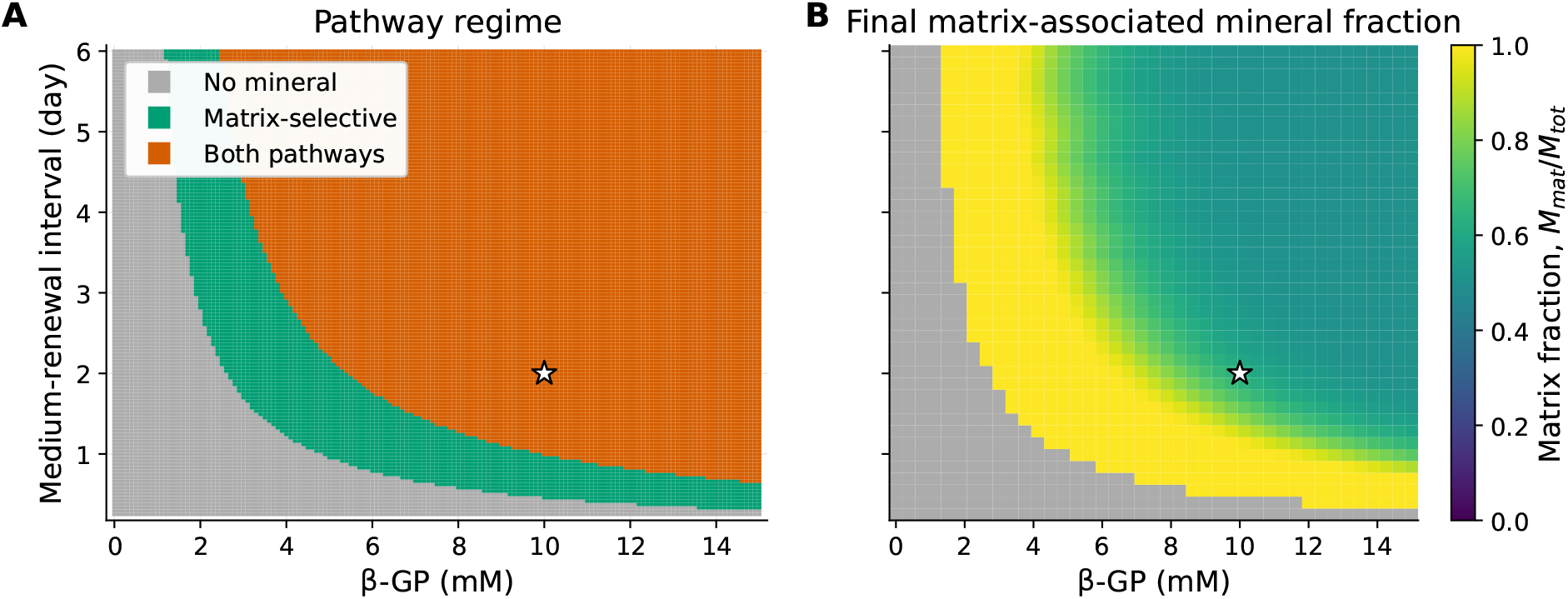
Analytical pathway-accessibility approximation and simulated endpoint partitioning. **A**, Classification from analytical crossing times assuming constant calcium and no mineral uptake of phosphate. The “Both pathways” category does not guarantee solution-driven activation in the coupled model. **B**, Day-28 matrix-associated mineral fraction, (*A* + *H*)*/*(*A* + *H* + *Q*), from the full coupled simulations. Dark regions in panel B indicate total retained mineral at or below 10^*−*3^ mM P-eq.; the dark category in panel A indicates that the analytical matrix threshold is not reached before renewal. The star marks the reference protocol.

To quantify the relative contribution of the matrix-associated pathway, we define the matrix-associated fraction in Eq. 10 as

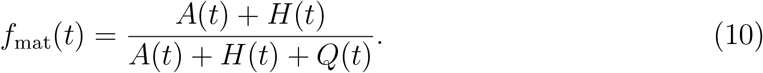

When both pathways were active, the final value *f*_mat_(*t*_*f*_ ) remained near or above 0.5 across the simulated domain (Fig. 3B). Across the endpoint grid, the minimum matrix-associated fraction was 0.50. This reflects the finite capacities used in the baseline parameter set: matrix-associated mineral approached *M*_mat,max_ = 1.0 mM P-eq. while solution-driven mineral approached *Q*_max_ = 1.0 mM P-eq., with a maximum simulated value of 1.00 mM P-eq. under high-supply conditions. The baseline protocol lies in the region where both pathways contribute, with a final matrix-associated fraction of ≈ 0.63.

### 4.3 Parameter sweeps identify supply and capacity controls

One-at-a-time sweeps showed that matrix-associated mineral accumulation was controlled most clearly by phosphate supply, hydrolysis efficiency, and matrix-associated capacity (Fig. 4). Without *β*-GP, no retained mineral accumulated. Increasing *β*-GP from 2.5 to 10 mM increased final *M*_mat_ from 0.00 to 1.00 mM P-eq.. Further increases to 15 and 20 mM produced only small additional increases in *M*_mat_, which approached the prescribed matrix-associated capacity, but they increased solution-driven mineral toward *Q*_max_.

**Figure 4.**
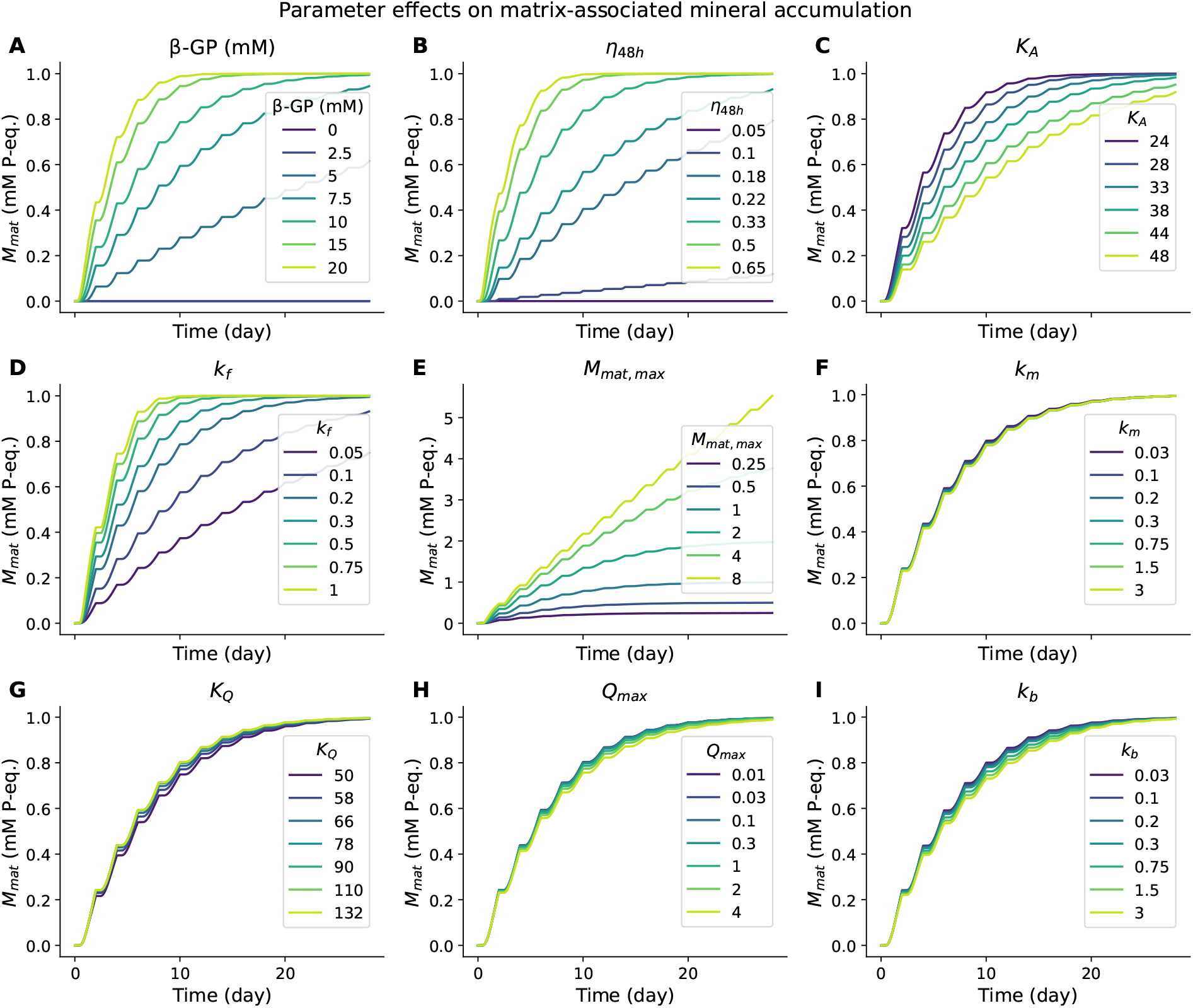
Dynamic parameter sweeps. Curves show *M*_mat_ = *A* + *H* over time while varying **A**, *β*-GP concentration; **B**, hydrolysis fraction *η*_48*h*_; **C**, matrix-associated onset parameter *K*_*A*_; **D**, matrix-associated formation coefficient *k*_*f*_ ; **E**, matrix-associated capacity *M*_mat,max_; **F**, maturation coefficient *k*_*m*_; **G**, solution-driven onset parameter *K*_*Q*_; **H**, solution-driven capacity *Q*_max_; and **I**, solution-driven formation coefficient *k*_*b*_. All other parameters were held at baseline values.

Increasing hydrolysis efficiency increased day-28 mineral retention. At *η*_48*h*_ = 0.05, neither pathway accumulated mineral. At *η*_48*h*_ = 0.33, *M*_mat_ = 0.998 and *Q* = 0.739 mM P-eq.; at 0.65, both approached 1.00 mM P-eq..

The relative separation of the pathway-onset thresholds strongly affected mineral partitioning. As *K*_*Q*_*/K*_*A*_ increased from 1.1 to 4.0, the onset of solution-driven precipitation was progressively delayed and the final matrix-associated fraction increased from 0.50 to 1.00, whereas final *M*_mat_ remained close to 1.00 mM P-eq. across the tested range.

Changing the two retention capacities had distinct effects on the final mineral distribution. Increasing *M*_mat,max_ from 0.25 to 8.00 mM P-eq. increased final *M*_mat_ from 0.25 to 5.53 mM P-eq., while reducing the relative solution-driven contribution. In contrast, increasing *Q*_max_ from 0.01 to 4.00 mM P-eq. increased final *Q* from 0.0067 to 1.62 mM P-eq. and lowered the final matrix-associated fraction from 0.99 to 0.38. The maturation co-efficient *k*_*m*_ mainly affected matrix-associated material from *A* to *H*: final *H* increased from 0.47 to 0.99 mM P-eq. as *k*_*m*_ increased from 0.03 to 3.00 day^*−*1^, while *M*_mat_ remained close to 1.00 mM P-eq..

### 4.4 Sensitivity of matrix-associated and solution-driven mineralization

Global sensitivity analysis separated the controls on final matrix-associated mineral from those on final solution-driven mineral (Fig. 5). Final *M*_mat_ was dominated by *M*_mat,max_, with a total Sobol index *S*_*T*_ = 0.74. The next largest effects were smaller: *η*_48*h*_ had *S*_*T*_ = 0.20, *β*-GP had *S*_*T*_ = 0.15, *K*_*A*_ had *S*_*T*_ = 0.05, and *k*_*f*_ had *S*_*T*_ = 0.03.

**Figure 5.**
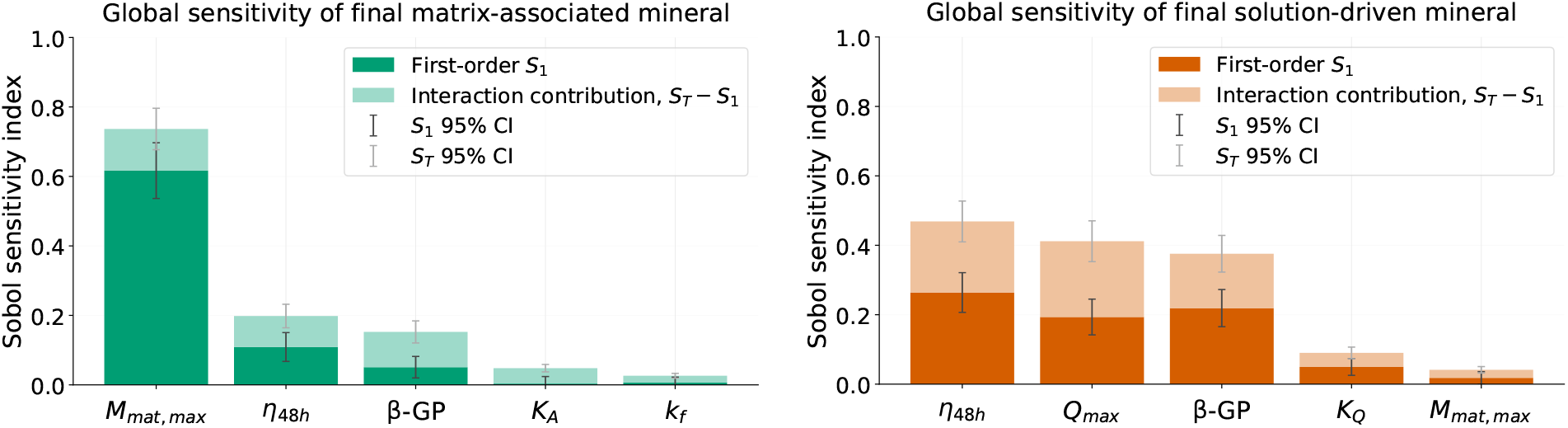
Global sensitivity of final pathway-specific mineral. **A**, Top five total Sobol indices for final matrix-associated mineral, *M*_mat_ = *A* + *H*. **B**, Top five total Sobol indices for final solution-driven mineral, *Q*. Bars separate first-order effects from interaction contributions.

The solution-driven endpoint had a broader control structure. Final *Q* was most sensitive to *η*_48*h*_ (*S*_*T*_ = 0.47), *Q*_max_ (*S*_*T*_ = 0.41), and *β*-GP (*S*_*T*_ = 0.38). *K*_*Q*_ also contributed (*S*_*T*_ = 0.09). *M*_mat,max_ had a smaller indirect effect (*S*_*T*_ = 0.04) under the tested parameter ranges.

### 4.5 How well do day-28 mineral results constrain the model?

We examined whether day-28 total mineral constrains pathway origin and parameter values (Fig. 6). An initial search of 2000 Latin-hypercube samples was followed by bounded optimization of the 50 closest candidates. Together, these searches identified 80 distinct parameter sets within 2% of the reference endpoint.

**Figure 6.**
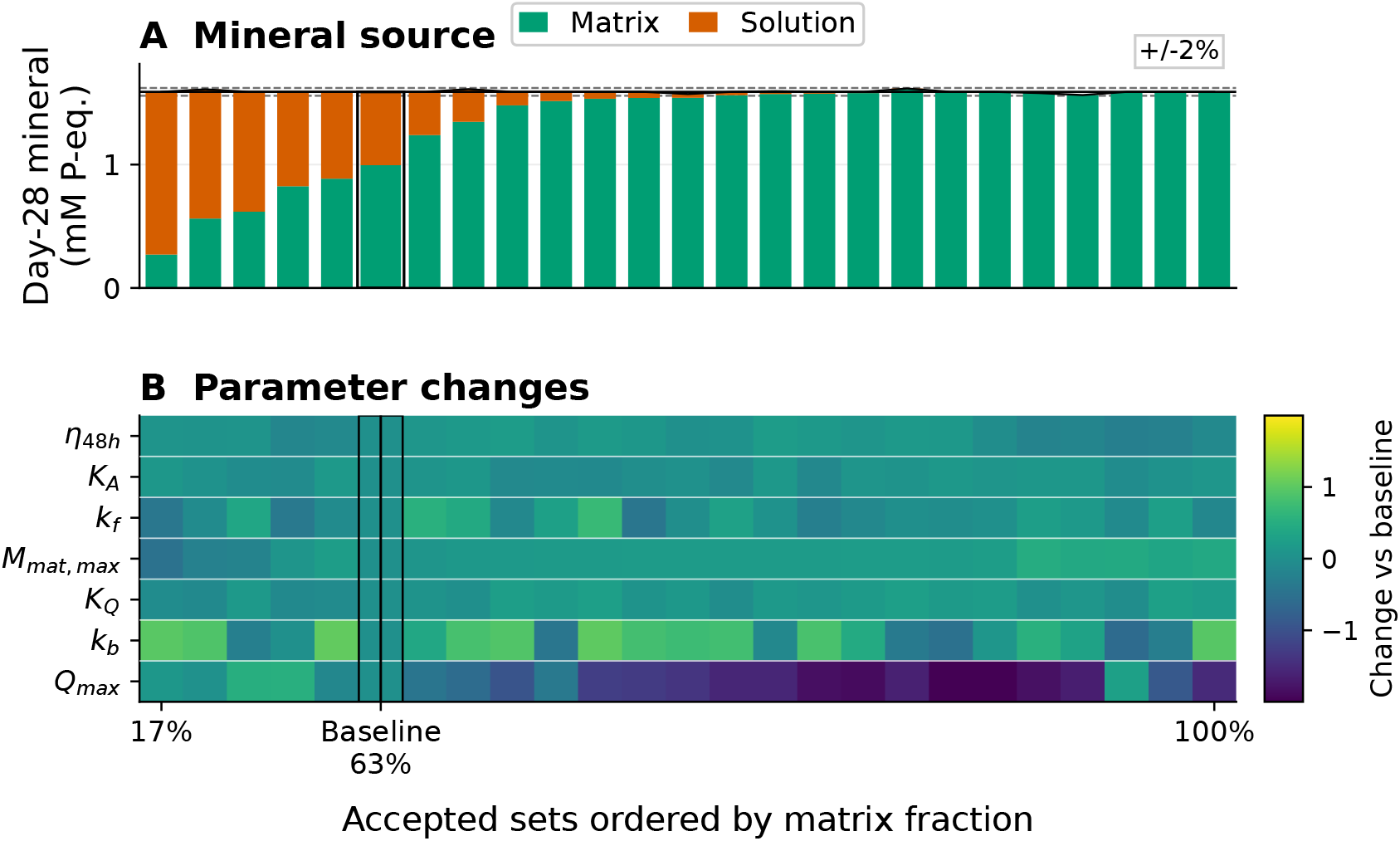
Day-28 endpoint equifinality. Of 80 parameter sets within 2% of the baseline total mineral, 24 are shown alongside the baseline. **A**, Matrix-associated and solution-driven contributions, ordered by matrix fraction. **B**, Corresponding parameter deviations from baseline, expressed as log_10_(*θ/θ*_0_).

Each accepted parameterization reproduced essentially the same terminal mineral end-point, with *M*_tot_(28) ranging from 1.56 to 1.62 mM P-eq. around the baseline value of

1.59 mM P-eq.. However, the underlying pathway partition varied widely. The matrix-associated fraction *f*_mat_(28) ranged from 0.17 to 1.00, meaning that the same total mineral signal could correspond to predominantly solution-driven retention in one case and entirely matrix-associated retention in another.

The accepted parameterizations also showed distinct parameter fingerprints. The largest deviations from baseline occurred mainly in *Q*_max_, *k*_*b*_, *M*_mat,max_, and *k*_*f*_, whereas *η*_48*h*_, *K*_*A*_, and *K*_*Q*_ varied over narrower ranges. Thus, a single terminal mineral measurement does not uniquely identify either the pathway contribution or the parameter combination responsible for the measured endpoint.

### 4.6 Comparison with literature data

The model was compared illustratively with literature mineralization time courses; no kinetic parameter was fitted to either comparison series (Fig. 7). For the Alizarin Red comparison, total retained P-equivalent mineral was scaled to the day-28 value of the comparison series [42]. The model increased from 8.81 at day 4 to 30.0 at day 28, compared with values of 2, 4, 5, 12, 15, and 30 entered in the comparison dataset. The model overpredicted early and intermediate signals, and final-point agreement was imposed by scaling. Identifying an ARS signal with total P-equivalent mineral assumes a common effective assay response across mineral pools; retained calcium would instead be represented by *r*_*A*_*A* + *r*_*H*_ (*H* + *Q*).

**Figure 7.**
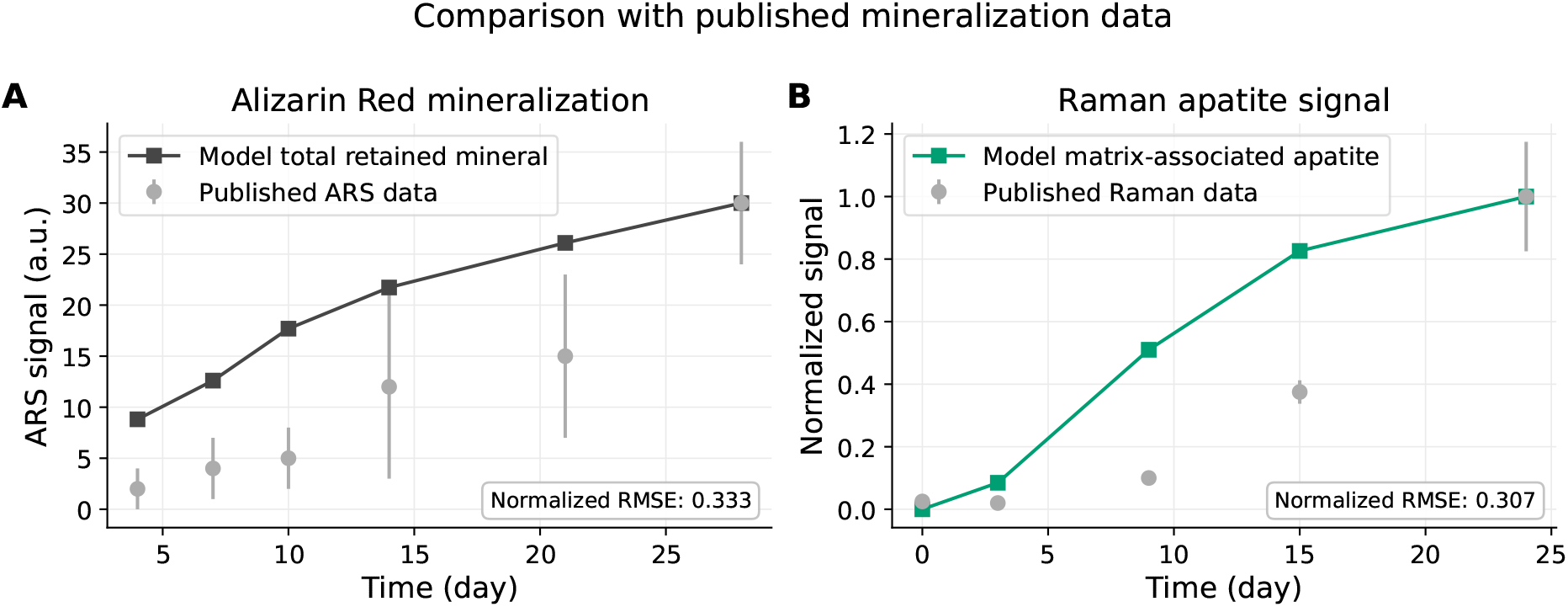
External consistency check against published mineralization time courses. **A**, Published Alizarin Red S data and day-28-scaled model total retained mineral. **B**, Published Raman apatite signal and day-24-normalized model matrix-associated apatite *H*(*t*). Corresponding RMSE values are given in the bottom right corner.

For the Raman comparison, the matrix-associated apatite-like pool *H*(*t*) was normalized to its day-24 value and simulated using 3-day medium renewal [43]. The normalized model values were 0.00, 0.08, 0.51, 0.83, and 1.00 at days 0, 3, 9, 15, and 24, compared with 0.025, 0.020, 0.100, 0.375, and 1.000 in the comparison series.

For Alizarin Red S, the normalized RMSE was 0.333 across five time points (days 4–21), excluding the day-28 endpoint used for scaling. For Raman, the RMSE was 0.307 across four time points (days 0–15), excluding day 24. Both comparisons showed a similar overall trend, although the model predicted faster mineral accumulation at intermediate times. This partly reflects our model structure, which does not account for ascorbic acid-dependent matrix formation or changes in mineralization capacity over time. Protocol-specific calibration would be needed for quantitative agreement.

### 4.7 Testable scenarios

The model suggests three experimentally testable predictions under the considered culture conditions. First, the ACP-like precursor is transient: it accumulates before apatite-like matrix mineral and subsequently declines as maturation proceeds. This sequence could be examined using time-resolved Raman, FTIR, or electron microscopy coupled with elemental analysis [44, 43, 45, 46].

Second, once matrix-associated mineral approaches its finite capacity, additional *β*-GP increases mainly solution-driven deposition, allowing bulk mineral accumulation to increase without a corresponding increase in matrix-associated mineral. This prediction could be tested using a *β*-GP dose series combined with spatially resolved mineral characterization [27, 19].

Third, shorter medium-renewal intervals are predicted to reduce phosphate accumulation and thereby delay or suppress solution-driven precipitation relative to matrix-associated mineralization. This could be examined by varying renewal frequency while monitoring soluble phosphate and the spatial distribution of mineral deposits [18, 22, 20].

## 5 Discussion

The model distinguishes matrix-associated mineralization from solution-driven calcium-phosphate deposition through two effective nucleation thresholds. By varying calcium and phosphate concentrations, *β*-GP concentration, hydrolysis efficiency, and medium-renewal frequency, the model provide a tool to examine how culture conditions influence the timing and extent of mineral formation through each pathway. These simulations can help narrow the experimental conditions to those expected to support matrix-associated mineralization with limited non-matrix deposition.

Under the reference condition (10 mM *β*-GP, *η*_48h_ = 0.30, 2-day renewal), about 3.0 mM phosphate was released per cycle. The matrix-associated criterion was reached at 11 hours, before the solution-driven criterion at 29 hours. This ordering is imposed through *K*_*A*_ *< K*_*Q*_, but is consistent with a lower effective nucleation barrier in matrix-supported environments [18, 9, 2]. Continued phosphate accumulation then allowed solution-driven deposition to increase further [27, 19, 21]. These threshold crossings indicate the onset of the modeled pathways, not necessarily detectable mineralization. Amorphous or crystalline mineral must accumulate sufficiently to produce a measurable signal.

The matrix-associated route also produced a transient ACP-like precursor. *A* peaked at ≈ 0.34 mM P-eq. on day 6 and then declined as *H* accumulated. This sequence is consistent with reports of amorphous or poorly ordered calcium-phosphate phases preceding more mature apatite [12, 13, 11, 6]. In the model, however, *A* should be interpreted as an effective precursor pool rather than a uniquely defined mineral phase.

The effect of phosphate supply was clearest at the endpoint. Increasing *β*-GP from 5 to 10 mM raised final *M*_mat_ from ≈ 0.62 to ≈ 0.99 mM P-eq., while *Q* increased from 0.00 to ≈ 0.59 mM P-eq.. Once *A* + *H* approached *M*_mat,max_, additional phosphate contributed mainly to the solution-driven pool. Thus, within the model, stronger endpoint staining at high phosphate supply does not necessarily indicate greater matrix-associated mineralization.

The regime map therefore suggests a practical matrix-selective window at intermediate *β*-GP supply and relatively frequent medium renewal. Renewal every 2–3 days combined with lower *β*-GP concentrations reduced the solution-driven contribution while preserving substantial matrix-associated mineralization. This partly matches primary osteoblast work, where 2–5 mM *β*-GP gave selective matrix mineralization and 10 mM gave widespread non-specific deposition [28].

Final *M*_mat_ was dominated by *M*_mat,max_ (*S*_*T*_ ≈ 0.74), whereas final *Q* was more sensitive to *η*_48h_, *Q*_max_, and *β*-GP supply (*S*_*T*_ ≈ 0.47, 0.41, and 0.38). More importantly, 80 parameter sets reproduced the day-28 total mineral within 2% of the reference endpoint, despite matrix-associated fractions ranging from 0.17 to 1.00, revealing the potential ambiguity in interpreting experimental results.

Biological variability can shift the effective model parameters. Differences between cell sources or subclones in matrix production, matrix-vesicle activity, and mineralization competence may alter both the onset and capacity of the matrix-associated pathway [10, 23]. A single parameter set should therefore not be expected to transfer directly between culture systems.

Calibration would benefit most from measurements that retain temporal or spatial information. Soluble calcium and phosphate between medium renewals could constrain phosphate release and pathway activation, while Raman or FTIR mapping could follow the transition from precursor-rich to mature mineral [44, 43, 45, 46]. Electron microscopy, diffraction, and targeted perturbations of collagen deposition, alkaline phosphatase activity, cell density, or matrix-vesicle abundance could further test mineral structure and pathway origin [1, 10, 2].

Several limitations remain. The model does not explicitly resolve aqueous speciation, spatial transport, or molecular nucleation mechanisms. Recent real-time studies of collagen mineralization further illustrate the difficulty of distinguishing mineral formation pathways experimentally [47]. The predicted pathway contributions therefore reflect the assumed model mechanisms rather than directly measured fractions. The model is intended to clarify how culture conditions influence mineral deposition, rather than predict mineral chemistry. Future work could incorporate time-dependent matrix capacity and alkaline-phosphatase activity, with dose-response, acellular-control, and time-course experiments providing additional constraints on pathway contributions.

## 6 Conclusion

The model illustrates how matrix-associated mineralization and solution-driven calcium-phosphate precipitation can affect the final mineral amount in osteogenic culture. Under the reference conditions, matrix-associated formation preceded solution-driven precipitation, while increasing phosphate supply mainly promoted non-matrix deposition once the assumed matrix capacity was approached. More importantly, parameter sets producing nearly identical day-28 mineral amounts showed substantially different pathway contributions, with matrix-associated fractions ranging from 0.17 to 1.00. This indicates that total mineral accumulation alone may provide limited information about the underlying deposition pathways. Although the relative contributions depend on the model assumptions and parameter values, the results reveal a potential ambiguity in interpreting conventional mineralization amounts. Measurements that distinguish mineral location and composition would help constrain these contributions and provide a basis for testing the proposed mechanisms experimentally.

## Funding and acknowledgments

This work was supported by EU Horizon 2020 (Marie Skłodowska-Curie grant No. 945371).

## Author Contributions

AAD: conceptualization, methodology, software, investigation, formal analysis, visualization, writing - original draft. TC: investigation, writing - review & editing.

## Declaration of Generative AI Use

Generative AI tools were used to assist with language editing, code development, and debugging. The mathematical model, numerical results, and scientific interpretations were reviewed and verified by the authors. The authors take full responsibility for the manuscript’s content.

## Data availability

Source code is publicly available on github.com/aliaslandemir/mineral-pathways.

