## Supplementary Information for "Distinguishing matrix-associated mineralization from solution-driven calcium phosphate precipitation in osteogenic culture"

This Supplement documents the numerical settings, parameter ranges, sensitivity and equifinality analyses, and numerical verification.

#### S1 Numerical settings

Simulations covered 28 days, with complete medium renewal every 2 days. The equations were integrated between renewals using RK45 (relative and absolute tolerances $10^{-6}$ and $10^{-9}$), with output every 0.05 day.

Table S1: Reference parameter values used for the deterministic simulation.

| Parameter | Value | Unit | Definition |
|---|---|---|---|
| $C_{\text{ext}}$ | 2.00 | mM | Fresh-medium soluble calcium |
| $P_{\text{ext}}$ | 1.25 | mM | Fresh-medium inorganic phosphate |
| $\beta$-GP | 10.0 | mM | Fresh-medium $\beta$-glycerophosphate |
| $\eta_{48\text{h}}$ | 0.30 | – | Fraction hydrolysed over 48 h |
| $K_A$ | 33.0 | $\text{mM}^5$ | Matrix-associated onset parameter |
| $K_Q$ | 66.0 | $\text{mM}^5$ | Solution-driven onset parameter |
| $k_f$ | 0.20 | $\text{day}^{-1}$ | Precursor-formation coefficient |
| $M_{\text{mat,max}}$ | 1.0 | mM P-eq. | Matrix-associated capacity |
| $k_m$ | 0.20 | $\text{day}^{-1}$ | Precursor-maturation coefficient |
| $k_b$ | 0.20 | $\text{day}^{-1}$ | Solution-driven formation coefficient |
| $Q_{\max}$ | 1.0 | mM P-eq. | Solution-origin retention capacity |
| $r_A, r_H$ | 1.245, 1.585 | – | Reference effective Ca/P ratios |

For the 100-run stochastic ensemble (seed 42), effective Ca/P ratios were sampled uniformly from $r_A \in [1.00, 1.49]$ and $r_H \in [1.50, 1.67]$. The reference values in Table S1 are their midpoints.

#### S2 Reference-run output

Reference-run quantities are given in Table S2.

#### S3 One-parameter sweeps

One parameter at a time was varied around the reference configuration, using the values in Table S3.

Outputs included $A(28)$, $H(28)$, $Q(28)$, $A(28) + H(28)$, total retained mineral, the matrix-associated fraction, and maximum $S_A$.

Table S2: Selected numerical outputs from the reference simulation.

| Quantity | Value |
|---|---|
| First numerical $S_A = 1$ crossing | 0.456 day |
| First numerical $S_Q = 1$ crossing | 1.202 days |
| Time with $S_A > 1$ | 21.350 days |
| Time with $S_Q > 1$ | 13.150 days |
| Peak $A$ | 0.343 mM P-eq. |
| Time of peak $A$ | 6.0 days |
| Final $A$ | 0.032 mM P-eq. |
| Final $H$ | 0.963 mM P-eq. |
| Final $Q$ | 0.594 mM P-eq. |
| Final $A + H$ | 0.995 mM P-eq. |
| Final $A + H + Q$ | 1.589 mM P-eq. |
| Final matrix-associated fraction | 0.626 |

Table S3: Values used in the one-parameter sweeps.

| Parameter | Reference | Sweep values |
|---|---|---|
| $\beta$-GP | 10.0 | 0, 2.5, 5, 7.5, 10, 15, 20 |
| $\eta_{48\mathrm{h}}$ | 0.30 | 0.05, 0.10, 0.18, 0.22, 0.33, 0.50, 0.65 |
| $K_A$ | 33.0 | 24, 28, 33, 38, 44, 48 |
| $k_f$ | 0.20 | 0.05, 0.10, 0.20, 0.30, 0.50, 0.75, 1.00 |
| $M_{\mathrm{mat,max}}$ | 1.0 | 0.25, 0.5, 1, 2, 4, 8 |
| $k_m$ | 0.20 | 0.03, 0.10, 0.20, 0.30, 0.75, 1.50, 3.00 |
| $K_Q$ | 66.0 | 50, 58, 66, 78, 90, 110, 132 |
| $Q_{\max}$ | 1.0 | 0.01, 0.03, 0.10, 0.30, 1.0, 2.0, 4.0 |
| $k_b$ | 0.20 | 0.03, 0.10, 0.20, 0.30, 0.75, 1.50, 3.00 |

#### S4 Global sensitivity analysis

Global sensitivity was assessed for nine inputs using scrambled Sobol sampling ($N = 1024$, seed 42), without second-order indices. For $D = 9$, this required $N_{\text{eval}} = N(D + 2) = 11264$ model evaluations. The outputs were $M_{\text{mat}}(28) = A(28) + H(28)$ and $Q(28)$.

Table S4: Parameter ranges used for the global sensitivity analysis.

| Parameter | Reference | Range | Unit |
|---|---|---|---|
| $\beta$-GP | 10.0 | 5–20 | mM |
| $\eta_{48\text{h}}$ | 0.30 | 0.10–0.50 | – |
| $K_A$ | 33.0 | 24–48 | $\text{mM}^5$ |
| $K_Q$ | 66.0 | 50–132 | $\text{mM}^5$ |
| $k_f$ | 0.20 | 0.05–1.00 | $\text{day}^{-1}$ |
| $M_{\text{mat,max}}$ | 1.0 | 0.25–4.0 | mM P-eq. |
| $k_m$ | 0.20 | 0.03–3.0 | $\text{day}^{-1}$ |
| $k_b$ | 0.20 | 0.03–3.0 | $\text{day}^{-1}$ |
| $Q_{\text{max}}$ | 1.0 | 0.01–4.0 | mM P-eq. |

Table S5: Largest Sobol indices for the two terminal pathway outputs.

| Output | Parameter | $S_1$ | $S_T$ |
|---|---|---|---|
| $A + H$ | $M_{\text{mat,max}}$ | 0.617 | 0.737 |
| $A + H$ | $\eta_{48\text{h}}$ | 0.109 | 0.198 |
| $A + H$ | $\beta$-GP | 0.051 | 0.152 |
| $A + H$ | $K_A$ | 0.004 | 0.048 |
| $A + H$ | $k_f$ | 0.006 | 0.026 |
| $Q$ | $\eta_{48\text{h}}$ | 0.264 | 0.469 |
| $Q$ | $Q_{\text{max}}$ | 0.193 | 0.412 |
| $Q$ | $\beta$-GP | 0.219 | 0.376 |
| $Q$ | $K_Q$ | 0.050 | 0.090 |
| $Q$ | $M_{\text{mat,max}}$ | 0.018 | 0.041 |

#### S5 Conditional trajectory-sensitivity profiles

For each selected parameter, the total-mineral trajectory was compared with the reference at days 4, 7, 10, 14, 21, and 28 using the normalized error

$$E(\theta) = \frac{1}{n} \sum_i \left[ \frac{M_{\text{tot}}(t_i; \theta) - M_{\text{tot}}(t_i; \theta_0)}{\max\{M_{\text{tot}}(t_i; \theta_0), 0.05 M_{\text{tot}}(28; \theta_0)\}} \right]^2 . \tag{S1}$$

The profiled parameters were $k_f$, $k_m$, $k_b$, $Q_{\text{max}}$, and $M_{\text{mat,max}}$. The rate constants and $Q_{\text{max}}$ spanned $10^{-2}$–$10^1$; $M_{\text{mat,max}}$ spanned $10^{-1}$–$10^1$, on the configured logarithmic grid. These are sensitivity diagnostics, not fitted parameter estimates.

#### S6 Structural $K_Q/K_A$ sensitivity

Threshold separation was tested by varying $K_Q/K_A$ at fixed $K_A = 33.0$. Ratios were 1.1, 1.25, 1.5, 1.75, 2.0, 2.4, 3.0, and 4.0; the reference ratio was 2.0.

Table S6: Endpoint effects of changing the solution-to-matrix onset ratio.

| $K_Q/K_A$ | $t_A$ | $t_Q$ | $t_Q - t_A$ | $A + H$ | $Q$ | Matrix fraction |
|---|---|---|---|---|---|---|
| 1.10 | 0.456 | 0.518 | 0.062 | 0.991 | 0.980 | 0.503 |
| 1.25 | 0.456 | 0.612 | 0.156 | 0.992 | 0.952 | 0.510 |
| 1.50 | 0.456 | 0.776 | 0.320 | 0.993 | 0.866 | 0.534 |
| 1.75 | 0.456 | 0.963 | 0.507 | 0.994 | 0.740 | 0.573 |
| 2.00 | 0.456 | 1.202 | 0.746 | 0.995 | 0.594 | 0.626 |
| 2.40 | 0.456 | 3.634 | 3.178 | 0.996 | 0.367 | 0.731 |
| 3.00 | 0.456 | 7.941 | 7.485 | 0.996 | 0.135 | 0.881 |
| 4.00 | 0.456 | 17.978 | 17.522 | 0.997 | 0.003 | 0.997 |

### S7 Day-28 endpoint equifinality

Day-28 equifinality was examined for $\eta_{48\mathrm{h}}$, $K_A$, $K_Q$, $k_f$, $M_{\mathrm{mat,max}}$, $k_b$, and $Q_{\max}$. Latin-hypercube candidates were sampled in log space within the Sobol bounds (Table S4; seed 271828). Sets within 2% of the baseline total retained mineral were accepted.

The reference day-28 values are $A = 0.032$, $H = 0.963$, $A + H = 0.995$, $Q = 0.594$, and $M_{\mathrm{tot}} = 1.589$ mM P-eq., with a matrix-associated fraction of 0.626. Accepted parameterizations were stored; the plot places the baseline column at its actual fraction.

### S8 Numerical verification

Calcium- and phosphate-equivalent balances were checked between renewals. The reference-run maximum relative errors were $6.32 \times 10^{-16}$ (phosphate) and $4.01 \times 10^{-16}$ (calcium). Across the stochastic ensemble, the mean maximum errors were $1.45 \times 10^{-11}$ and $1.35 \times 10^{-10}$, respectively; the corresponding 97.5th percentiles were $4.67 \times 10^{-13}$ and $4.06 \times 10^{-12}$.

#### S8.1 Numerical step-size sensitivity

The reference simulation was repeated with maximum RK45 steps and ion-availability limiting intervals of 0.10, 0.05, 0.025, and 0.0125 day. Parameters, renewal conditions, and solver tolerances were unchanged.

The total-mineral trajectories nearly overlapped (Fig. S1A). Relative to the finest step, day-28 total mineral differed by at most 0.00203% and the matrix-associated fraction by at most 0.000438 percentage points (Fig. S1B).

The limiter was not activated in the reference runs or in 50 additional Latin-hypercube combinations spanning the nine sensitivity ranges. These checks apply to the tested conditions, not the full parameter space.

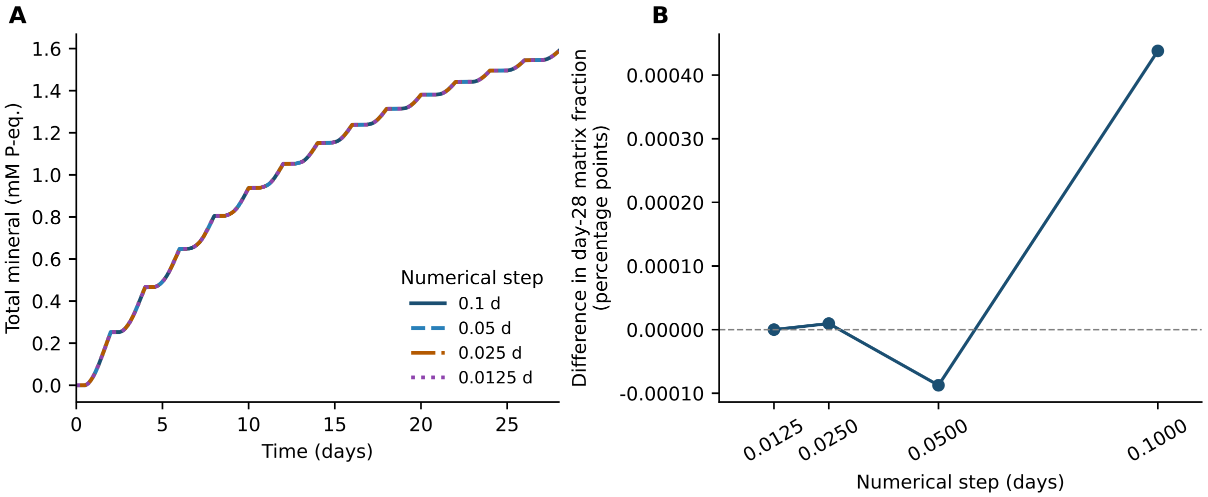


Figure S1: Numerical step-size sensitivity of the mineralization model. (A) Total retained mineral over 28 days for numerical step settings of 0.10, 0.05, 0.025, and 0.0125 day. (B) Difference in the day-28 matrix-associated mineral fraction relative to the finest-step simulation (0.0125 day), expressed in percentage points. Both the maximum RK45 step and the ion-availability limiting interval were varied, while all biological parameters and solver tolerances were unchanged.
